# Early alphavirus replication dynamics in single cells reveal a passive basis for superinfection exclusion

**DOI:** 10.1101/2020.09.07.282053

**Authors:** Zakary S. Singer, Pradeep M. Ambrose, Tal Danino, Charles M. Rice

**Affiliations:** Department of Biomedical Engineering, Columbia University, New York, NY 10027, USA; Laboratory of Virology and Infectious Disease, The Rockefeller University, New York, NY, 10065, USA; Herbert Irving Comprehensive Cancer Center, Columbia University, New York, NY 10027, USA; Data Science Institute, Columbia University, New York, NY 10027, USA

**Keywords:** Alphavirus, RNA virus, superinfection exclusion, single-cell, single-molecule, replication dynamics, smFISH, time-lapse

## Abstract

While decades of research have elucidated many steps in the alphavirus lifecycle, the earliest replication dynamics have remained unclear. This missing time window has obscured early replicase strand synthesis behavior and prevented elucidation of how the resulting activity gives rise to a superinfection exclusion environment, one of the fastest competitive phenotypes among viruses. Using quantitative live-cell and single-molecule imaging, we characterize the strand preferences of the viral replicase *in situ*, and measure protein kinetics in single cells over time. In this framework, we evaluate competition between alphaviruses, and uncover that early superinfection exclusion is actually not a binary and unidirectional process, but rather a graded and bidirectional viral interaction. In contrast to competition between other viruses, alphaviruses demonstrate a passive basis for superinfection exclusion, emphasizing the utility of analyzing viral kinetics within single cells.

## Introduction

Sindbis virus is a plus-strand RNA virus with a broad host range that cycles between vertebrate and mosquito hosts **(Strauss and Strauss 1994)**. As the type virus of the alphavirus genus, its replication has been studied in detail since its discovery in 1953 **(Taylor and Hurlbut 1953, Taylor, Hurlbut et al. 1955)**. Upon release into a cell, the infecting plus strand is translated as a large polyprotein that contains the replication machinery. Polyprotein processing through cleavage is tightly regulated, where differential cleavage between early and late infection alter the template preferences of the replicase. This progression has been suggested to cause a transition from its complementary minusstrand production, then to plus-strand replication, and finally to sub-genomic transcription, the last of which encodes the viral structural proteins **(Groot, Hardy et al. 1990, Sawicki, Barkhimer et al. 1990, Lemm and Rice 1993, Lemm and Rice 1993, Sawicki and Sawicki 1993, Lemm, Rümenapf et al. 1994, Shirako and Strauss 1994)**.

Across both prokaryotes and eukaryotes, a phenomenon know as superinfection exclusion (SE) has been observed, where infection by one virus can block replication of a subsequent homologous virus **(Dulbecco 1952, Steck and Rubin 1966, Bratt and Rubin 1968, Whitaker-Dowling, Ungner et al. 1983, Johnson and Spear 1989, Lee, Tscherne et al. 2005, Zou, Zhang et al. 2009, Biryukov and Meyers 2018)**. In principle, such a behavior could serve to improve dissemination by preventing reinfection of already infected cells, and protect a virus from competing with its progeny or a similar superinfecting virus. One potential mechanism of action is limiting a second virus’ entry, through down-regulation or depletion of receptors, or even repulsion of virions away from the surface of infected cells **(Garcia and Miller 1991, Huang, Li et al. 2008, Stiles, Milne et al. 2008, Doceul, Hollinshead et al. 2010)**. The changes required in cellular state to achieve this are generally thought to occur over the course of several hours following infection. However, for alphaviruses like Sindbis virus, such a phenomenon can be seen within just 15 minutes of the first infection, leading to reduction in titer of the second alphavirus by an order of magnitude **(Johnston, Wan et al. 1974, Adams and Brown 1985, Karpf, Lenches et al. 1997, Singh, Suomalainen et al. 1997)**.

Even though this strikingly rapid competitive behavior was observed for Sindbis over 40 years ago, its origins have remained elusive. Moreover, despite elegant studies revealing the lifecycle of alphaviruses, our understanding of the very earliest stages of replication and the specific events establishing such a rapid exclusionary environment are unclear. For example, population-level measurements have revealed the accumulation of plus and minus-strand RNA and concomitant processing of viral polyprotein over the first ~3-6 hours of infection, and the first release of progeny virions as early as ~4hpi **(Sawicki and Sawicki 1980, Sawicki, Sawicki et al. 1981, Strauss and Strauss 1994)**. However, it is unknown when the very first minus strand is produced, whether the early replicase prefers to synthesize plus or minus strand RNA within the first few hours of replication, and how such early replicase behaviors might achieve superinfection exclusion.

A major challenge in the study of early replication dynamics and how theses behaviors might influence competitive interactions has been the use of population-based measurements that lack the sensitivity to interrogate low-abundance targets during early infection. While classic studies have shown the average growth of the virus over time across millions of cells, what are the underlying growth kinetics that contribute to this mean, and how might the inherent cell-to-cell variability and unsynchronized nature of a spreading infection obscure these measurements? For example, does the virus grow constantly and linearly in a subpopulation, or grow exponentially and then plateau in some cells and decay over time in others? A quantitative analysis of early replicase activity, and the ability to follow the time course in single-cells could elucidate the underlying nature of viral replication, and reveal events on the time scale during which superinfection exclusion behaviors emerge (**Fig. 1**).

**Figure 1).**
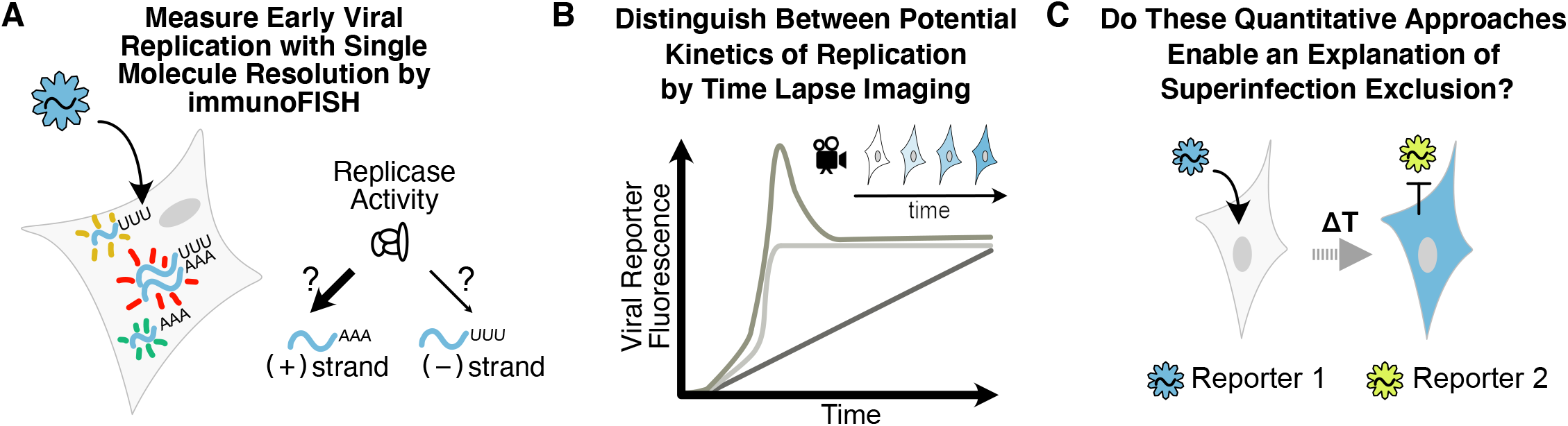
Using quantitative single-cell measurements to uncover replication kinetics of Sindbis virus. **A)** Simultaneous measurements of early vRNA species by immuno-FISH can reveal early replicase preferences in the production of plus-strand or minus-strand RNA, and the trajectory of distinct vRNA species (plus, minus, duplex) strands over time in distributions of fixed cells. **B)** Following viral replication in live single cells over time can uncover the temporal changes in viral kinetics within the same cell, and distinguish between possible modes of viral growth. **C)** Through these single-cell studies of viral replication, can the observed behaviors be used better understand early viral competition in the superinfection exclusion phenomenon?

The use of single-cell analyses in biology have enabled an unprecedented look into the behavior of natural genetic circuits, in part, by revealing how the variability of individual cells can be masked by an overall population’s behavior **(Fraser and Kærn 2009, Locke and Elowitz 2009, Raj and van Oudenaarden 2009, Chalancon, Ravarani et al. 2012)**. The use of single-cell approaches in virology has only recently begun to show how variability between individual cells contribute to viral growth and spreading kinetics; how stochasticity can influence fate-selection in both phages and animal viruses; and how quantitative models might lead to improved antivirals **(Snijder, Sacher et al. 2009, Doceul, Hollinshead et al. 2010, Jones, Catanese et al. 2010, Zeng, Skinner et al. 2010, Chou, Vafabakhsh et al. 2012, Schulte and Andino 2014, Heldt, Kupke et al. 2015, Razooky, Pai et al. 2015, Akpinar, Timm et al. 2016, Ramanan, Trehan et al. 2016, Guo, Li et al. 2017, Hansen, Wen et al. 2018, Drayman, Patel et al. 2019, Shnayder, Nachshon et al. 2020).** In the present work, by directly exploring the progression of viral replicase activity *in situ* and the resulting single-cell replication kinetics, we shed light on classic questions remaining in alphavirology, and provide a new framework for understanding early replication and the resulting exclusionary phenomenon.

## Results

### Strand-specific single molecule measurements during early viral RNA synthesis

In order to capture the dynamics of the earliest stages of replication, it is necessary to utilize an approach with high enough sensitivity to measure, simultaneously, individual molecules of multiple viral RNA species at low-abundance. Furthermore, the methods must permit interrogation at the single cell level, in order for the resulting measurements to reveal potential heterogeneity among the population, such as a refractory subset of cells, or an unsynchronized infection from spatial and temporal waves of spreading. Single molecule RNA fluorescence *in situ* hybridization (smFISH) can yield an assay in which the binding of multiple fluorescent probes to their target RNA creates discrete and countable diffraction limited spots (**Fig. 2A**). Furthermore, multiple chromatically distinct probe sets can be used in parallel to measure, with strand specificity, the abundance of the full length genomic plus-strand and its template, the minus-strand. Based on the targeting of these probe sets to the sequence of the non-structural proteins, levels of subgenomic RNA will not contribute to their signal. However, viral plus and minus strands are entirely complementary to each other and potentially form a duplex that cannot be bound by small 20mer smFISH probes under standard hybridization conditions. Therefore, we additionally perform immunostaining in the same sample using an antibody that recognizes stretches of duplex, or double-stranded RNA (dsRNA), longer than 40 base pairs **(Schonborn, Oberstraβ et al. 1991)**. Together, with this immunoFISH approach, we are able to distinguish free plus- and minus-strands from the fraction of viral RNA (vRNA) in duplex form, using automated image analysis to detect and quantify these individual puncta within segmented cells (**Fig. 2B**).

**Figure 2).**
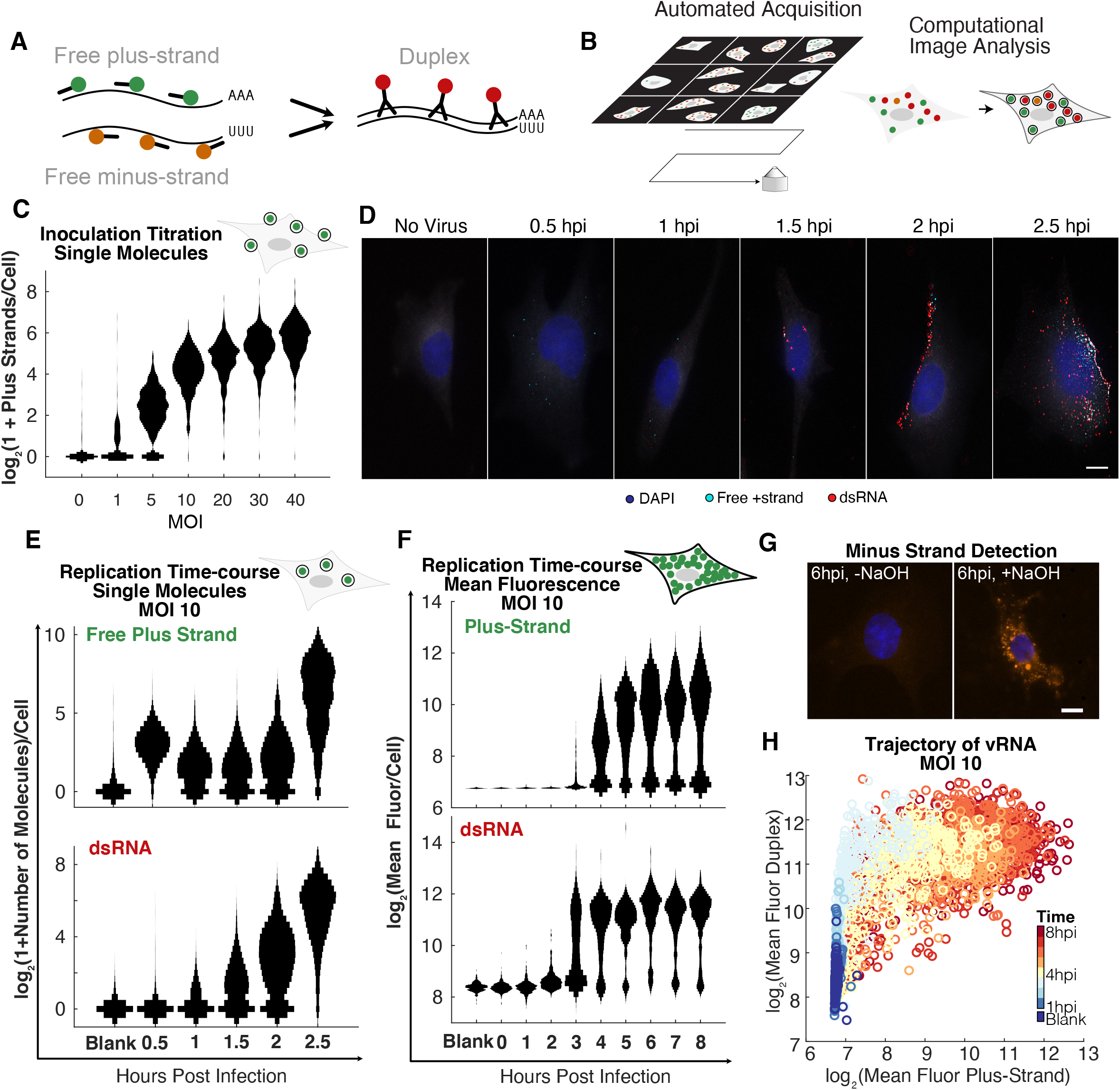
Trajectories of Sindbis vRNA by strand specific immunoFISH. **A)** Schematic showing the immunoFISH strand specific probes against free plus- and minus strands, as well as immunostaining of dsRNA by J2 antibody, each chromatically distinct from one another. **B)** Grid-based acquisition is performed automatically on the microscope (left), followed by an automated cell-segmentation and dot detection analysis (right). **C)** Counts of single spots of virions per cell immediately following inoculation, over a range of MOIs, displayed as smoothened vertical histograms. **D)** Representative images of infected cells, co-stained for dsRNA and free plus-strand RNA through 2.5hpi. Blue, DAPI; white, cytoplasm; cyan, free-plus strand; red, dsRNA. Scale bar is 10um. **E)** Counts of spots in individual cells of free plus-strand (top), and dsRNA (bottom) over a time course through 2.5h, displayed as smoothened vertical histograms. **F)** Distributions of total fluorescence per cell from free plus-strand (top) and dsRNA (bottom) over an 8 hour time course. **G)** Minus-strand RNA only appears after treatment with NaOH (right) and not without (left), shown at 6hpi. Scale bar is 10um. **H)** Single cell scatter plot of total fluorescence per cell for dsRNA vs free plus-strand RNA, where the color represents the sample’s time-point.

To establish the sensitivity of the method for detecting viral plus-strands, we exposed BHK-21 cells to virus across a gradient of MOIs and fixed the cells immediately following a 30 minute inoculation (**Fig. 2C, S1A**). At MOI 1, about half of all cells had detectable puncta of viral genomic RNA, with an average of 3.0±3.7 per cell, while at MOI 10, nearly all cells had puncta, with an average of 19.9±12.6 (mean±SD). These measurements also revealed a surprising amount of variability between cells. Measures of dispersion such as the Fano factor (variance/mean) provide a way to compare variability between distributions. For a Poisson distribution, the Fano factor is 1. At MOI 10, the observed Fano factor is 7.9=1±0.8 (mean±SEM), suggesting a dispersion greater than would be expected from a Poisson distribution **(Ellis and Delbrück 1939, Dulbecco 1952).** Together, these results confirm the sensitivity of the approach, and the ability to detect individual molecules of viral plus-strand RNA.

We next asked how the distributions of free plus- and minus-strands and duplex RNA evolved across the first 2.5 hours post infection (hpi), sampling cells at 30-minute intervals (N=3,553). Representative images are shown in **Figure 2D**. Intriguingly, the distribution of plus-strands detectable immediately following the inoculation (0.5 hpi) was significantly reduced at 1hpi (two-sample KS test, p=1.49×10^−67^), in conjunction with an increase in the first measurable duplex RNA (two-sample KS test, p=3.8×10^−16^**, Fig. 2E**). The emergence of duplex necessarily implies a corresponding production of minus-strand RNA from plus-strand, detectable as dsRNA. Furthermore, while dsRNA continues to increase steadily over this interval, free plus-strands did not. Between 2 and 2.5 hours, however, there was a sudden and significant increase in both dsRNA and free plus strands, and the abundance of plus-strand thereafter exceeded the limit of individually quantifiable puncta.

Broadening the window of investigation, we measured hourly time-points between 1hpi and 8hpi and quantified the resulting total fluorescence from plus-strand FISH and duplex RNA immunostaining in a total of 4,555 cells (**Fig. 2F**). Inspecting the temporal evolution of these distributions, a subpopulation of cells appeared to reach plateau levels of dsRNA as early as 3hpi, while the signal from free plus-strand production appeared to continue in most cells for about two additional hours. This upper-bound in dsRNA abundance is likely a result of a so called “minus-strand shutoff”, where the accumulated production of nsP2-protease leads to the cleavage of viral polyprotein into its constituent non-structural proteins, which form a replicase unable to produce additional minus-strands **(Sawicki and Sawicki 1980, Lemm, Rümenapf et al. 1994)**. Looking sub-cellularly, sites of dsRNA were concentrated most strongly along the boundary of the cell (**Fig. 2D**), consistent with a model in which replication occurs within spherules along the plasma membrane throughout the full time-course of infection **(Froshauer, Kartenbeck et al. 1988, Frolova, Gorchakov et al. 2010, Spuul, Balistreri et al. 2010)**. Strikingly, throughout these experiments, no free minus-strand RNA was observed (**Fig. S1B**). To rule out the possibility that the lack of detectable free minus-strand was caused by poor sensitivity of minus-strand probes, RNA duplexes disrupted through sodium hydroxide treatment were able to reveal minus-strand RNA (**Fig. 2G**).

As these dsRNA and plus-strand measurements are performed simultaneously in the same cells, we can additionally explore the evolution of the polarity of the replicase activity for either plus-strand or minus-strand (contained in duplex) over time. Plotting the single-cell measurements of total fluorescence of both species color-coded by time, a trajectory of viral replication emerges in which plus- and minus-strands are first produced in nearly equal amounts, yielding duplex RNA. Only after significant levels of duplex RNA are established in a given cell does free plus-strand then emerge (**Fig. 2H, S1C**).

Together, these single molecule and total fluorescence measurements provide one of the earliest detections of alphaviral replication, revealing the initiation of replication within the first hour of infection. Furthermore, these results suggest a revised model of early replication wherein both plus- and minus-strands are made at a similar rate during early infection - as opposed to a window that is biased towards minus strand synthesis - where both full-length viral RNAs can be utilized as templates during the first two hours of infection.

### Single-cell dynamics of viral replication

While smFISH provides a strand-specific view of replication at the level of the viral genome, the resulting kinetics are necessarily estimated from changes in distributions of cells taken at fixed time-points, and don’t consider the proteins responsible for the observed dynamics. Thus, the ability to follow replication in single cells over time and at the protein level would serve to complement these static RNA snapshots.

In order to evaluate the early replication dynamics of Sindbis virus in live cells, we inserted the fluorescent protein mTurquoise2 gene into the hyper variable region of the non-structural protein nsP3 (**Fig. 3A-B, Supplemental Movie 1**), which we designate as SINV/nsP3-mTurquoise2. The fluorescence from this fusion protein reporter provides a direct readout of non-structural protein levels, while having minimal effect on viral replication **(Jose, Taylor et al. 2017)**. To first establish the relationship between the fluorescent reporter and viral RNA, we quantified nsP3-mTurquoise2 levels simultaneously with viral RNA by immunoFISH. Reporter fluorescent kinetics were similar to viral RNA kinetics, plateauing at about 4hpi at MOI 10, with individual puncta observable robustly at 1.5hpi (**Fig. 3C, S2A**). Comparing protein levels to viral RNA, nsP3-mTurquoise2 appeared well correlated with dsRNA (Pearson correlation coefficient r=0.92, **Fig. 3D**), and just slightly less well correlated with free plus-strand (Pearson correlation coefficient r=0.90, **Fig. 3E**). Together, this data suggest that viral RNA and polyprotein levels increase with similar dynamics, and that following nsP3-mTurquoise2 over time can serve as a proxy for the underlying RNA levels as well.

**Figure 3).**
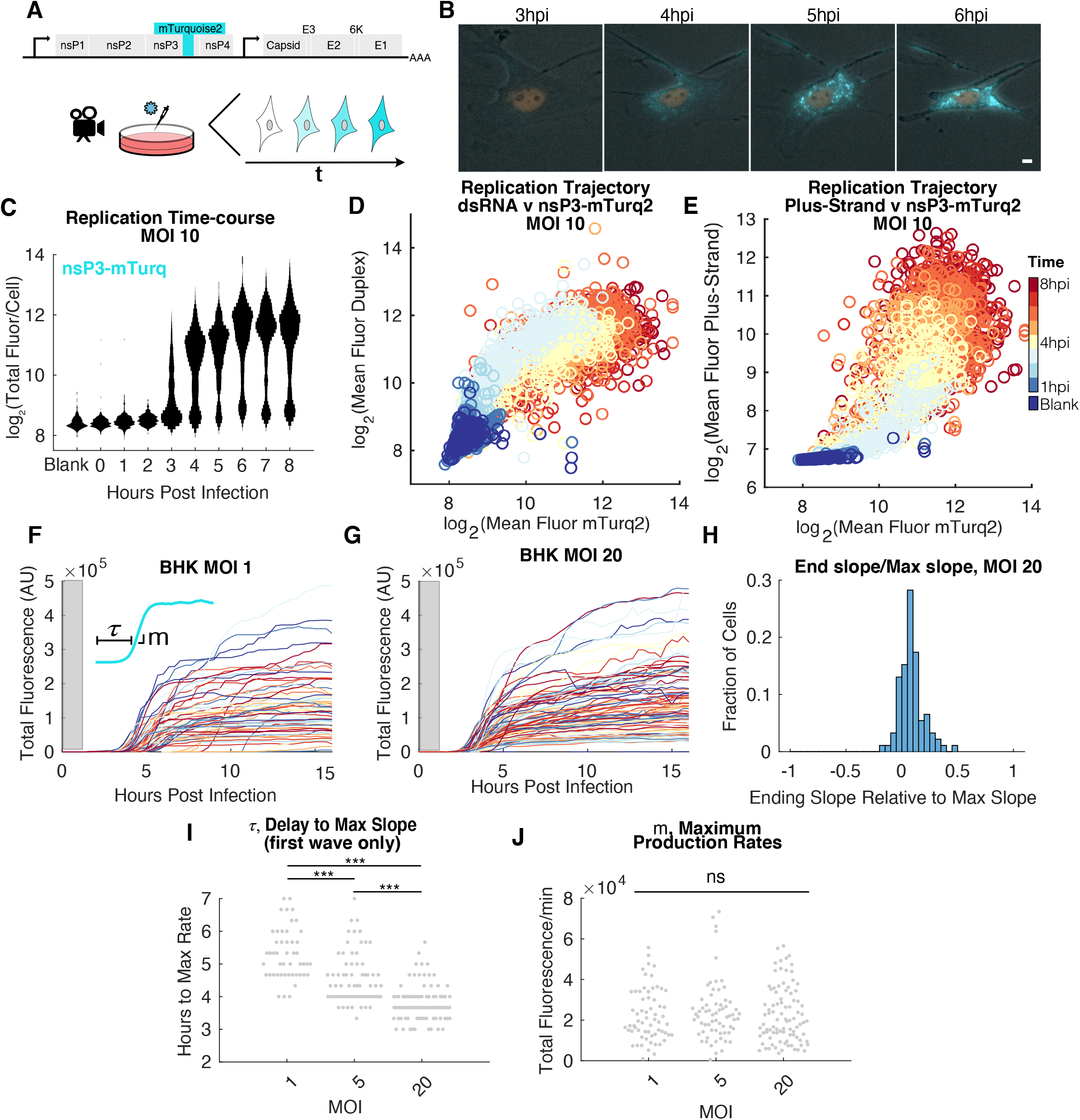
Time-lapse microscopy reveals logistic growth of viral replication. **A)** Schematic of viral reporter, and experimental approach. **B)** Example multicolor images of the same cell over time with a nuclear stain and increasing levels of nsP3-mTurquoise2. Scale bar is 10um. **C)** Single-cell distributions of total nsP3-mTurquoise2 normalized by area, plotted over time as vertical smoothened histograms. **D)** Scatter plot of total levels, normalized by area, of dsRNA vs nsp3-mTurquoise2, where each point is a single cell. **E)** Scatter plot of total levels, normalized by area, of plus-strand RNA vs nsp3-mTurquoise2, where each point is a single cell. **F)** Single cell traces of total fluorescence infected at MOI 1. Gray box indicates the first hour during the inoculation and movie-preparation window before measurements are taken. Inset indicates τ, the time delay until the trace’s maximum slope, *m*, on an example trace. **G)** Single cell traces of total fluorescence infected at MOI 20. Gray box indicates the first hour during the inoculation and moviepreparation window before measurements are taken. **H)** The fold reduction in slopes computed as the slope over the last three hours of the traces fit to a line, divided by the maximum growth of the trace, shown for MOI 20 infection. **I)** Distributions of the time delay τ, in hours, as a function of MOI, where each gray dot is a cell. Comparing MOI 1 to MOI 5 distributions, significance by the non-parametric two-sample Kolmogorov-Smirnov (KS) test is p=3.4×10^−7^. Comparing MOI 5 to MOI 20, p=5.5×10^−7^. Comparing MOI 1 to MOI 20, p=1.98×10^−19^. **J)** Distributions of the maximum production rates, *m*, as a function of MOI, where each gray dot is a cell. No comparisons are statistically significant by KS test.

To analyze replication within single cells in real-time, we adapted and expanded segmentation and tracking software based on nuclear fluorescence **(Hormoz, Singer et al. 2016)** to additionally follow cytoplasmic signals, where alphaviral replication occurs exclusively. This method yields a fluorescent “trace” for each cell that describes the temporal history of its nsP3-fused fluorescent reporter over the course of the experiment. We performed infections over a range of MOIs and asked whether and how replication dynamics differed between them. Strikingly, traces of infected cells revealed stereotyped logistic growth kinetics over all MOIs, where a sharp exponential increase in fluorescence was followed by a significantly reduced slope or a complete plateauing in production, similar to what was observed by immunoFISH (**Fig. 3F-G, S2B**). On average, the slope of the fluorescence over the last five hours of the movie was reduced 93±5% (mean±SD) relative to the maximal slope during early replication (**Fig. 3H, S2C**). Additionally, metrics quantifying trace parameters such as the delay to the onset of exponential growth designated as τ, and the maximum slope, *m*, can also be quantified, and compared between conditions. While logistic growth appeared consistent across these conditions, the onset delays, τ, appeared to decrease with increasing MOIs (**Fig. 3I**), while the maximum slope distributions appeared independent of MOI (**Fig. 3J**).

One potential explanation for these stereotyped kinetics is that a host-cell shut-off might lead to diminishing rates of replication or non-structural protein translation. To test if the kinetics are dominated by host cell transcription and translation shut-off, we utilized a double mutant shown to abrogate these functions through point mutations in the nsP2 protease domain and nsP3 macrodomain, respectively **(Akhrymuk, Frolov et al. 2018)**. Strikingly, in this mutant there is little to no change on the qualitative stereotyped kinetics, suggesting that the observed growth is not mediated by these two host effects (**Fig. S2D**).

### Superinfection exclusion arising from logistic growth

What are the implications of these apparent timescales and logistic growth, and how might it relate to observations of alphaviral competition? Previous work has shown that alphaviruses rapidly inhibit the replication of subsequent alphaviral infections in both mosquito and vertebrate cells, and is independent of interferon and defective interfering particles **(Zebovitz and Brown 1967, Stollar and Shenk 1973, Johnston, Wan et al. 1974, Igarashi, Koo et al. 1977, Adams and Brown 1985, Karpf, Lenches et al. 1997)**. For example, Sindbis virus has been shown to block infection by chikungunya, Una, Ross River, and Semliki Forest viruses, reducing their titer by between 2 and 5 orders of magnitude **(Eaton 1979, Karpf, Lenches et al. 1997)**. Such work, however, has necessarily relied on indirect measurements of replication provided by plaque assays, in which the progeny of the two competing viruses were distinguished by characteristic differences in temperature sensitivity, or by plaque size or shape. Thus, the relative competition and replication between the two viruses within the same cell at early times of replication have remained unclear. Though it has been previously shown that the inhibition to superinfection replication occurs post-entry **(Adams and Brown 1985)**, there has only been speculation about the potential sources of the blockade.

To examine some of the earliest events in the competition between two viruses in the same cell, we constructed a second reporter strain, replacing mTurquoise2 with the structurally nearly-identical but chromatically distinct mCitrine protein. Using these two reporters, we systematically varied the interval between inoculation of the two viruses, as well as the MOI of the second virus. Specifically, the first virus, SINV/nsP3-mTurquoise2 was inoculated at an MOI 10, and was introduced either 0, 20, 40, or 60 minutes before the second virus, SINV/nsP3-mCitrine. This second virus was added at a range of MOIs (1, 5, 10 or 30), after which we recorded the resulting fluorescence from both viruses using single-cell time-lapse imaging. An example multi-color image montage at 12hpi is shown in **Figure 4A**.

**Figure 4).**
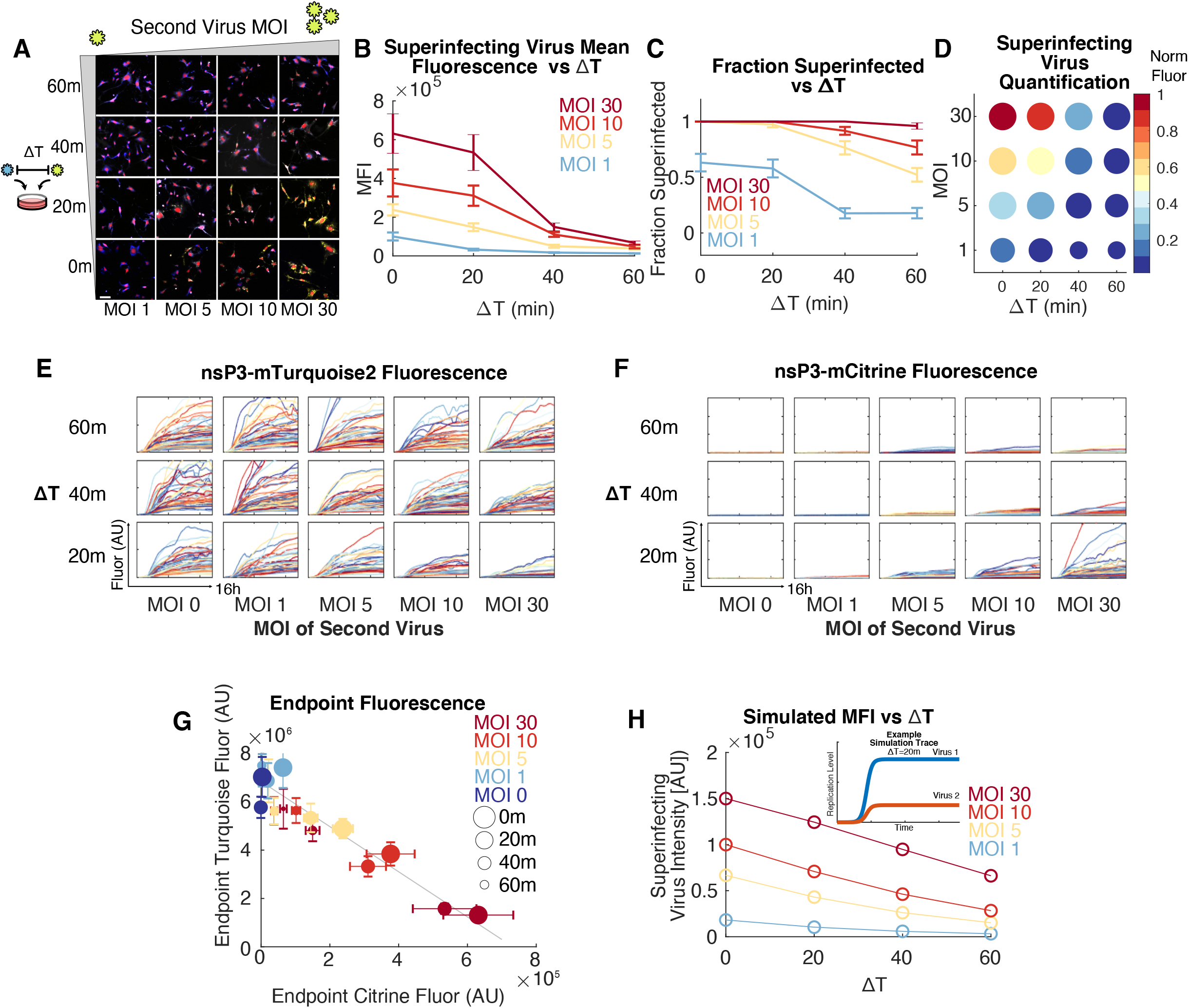
Single cell measurements of superinfection exclusion reveal bidirectional viral competition. **A)** Cells initially infected with MOI 10 of SINV/nsP3-mTurquoise2 are subsequently infected with SINV/nsP3-mCitrine at a range of MOIs (0, 5, 10, or 30) after different delays (either 0m, 20m, 40m, or 60m). Representative images show a nuclear stain (red), SINV/nsP3-mCitrine (yellow) and SINV/nsP3-mTurquoise2 (blue) under each condition described. Scale bar is 100um. **B)** Total fluorescence per area (or MFI) averaged over all cells in the given condition as a function of delay (ΔT), shown as mean±SEM. Color coding of each line represents a different superinfecting MOI. **C)** A plot of the fraction of cells infected by the second virus, SINV/nsP3-mCitrine as a function of delay (ΔT) and superinfecting MOI. Error bars are bootstrapped SEM. **D)** Information from B and C are combined to visually depict how temporal delays and relative MOIs together influence superinfection. Color represents the average normalized fluorescence intensity, while circle size represents the fraction of cells infected. **E-F)** Single cell traces of replication kinetics from mTurquoise2 (left) and mCitrine (right) reporter viruses under the experimental conditions described in A through the 13h time course. Y-axes of subplots in E are all equal, as are the Y-axes in F MOIs indicated are of the superinfecting SINV/nsP3-mCitrine. The first virus is always kept at MOI 10. **G)** Total fluorescence averaged over single-cells of nsP3-mTurquoise2 versus nsP3-mCitrine at the end of the superinfection movie (13hpi) shown in E and F Circle size represents delay between the first and second viruses, and color represents the MOI of the second virus, each shown at mean±SEM. Gray line is a linear-least squares fit, with a Pearson correlation coefficient of r=-0.99. **H)** Lotka-Volterra simulations of the superinfection experiment where both the first and second viruses are modeled identically (r=0.03/min and K=200,000, see Methods), over a range of delays and superinfecting MOIs. Inset shows a model of viral kinetics, where both begin at MOI 10, but with ΔT=20min.

Instead of displaying an all-or-nothing resistance to superinfection, infection by the second virus occurred in a graded manner dependent on the delay and relative MOI between the two viruses: as the delay between the two increased, or the MOI of the second virus was reduced, the ability of the second virus to replicate became further inhibited. This was reflected in both the mean fluorescence of the second virus SINV/nsP3-mCitrine in doubly infected cells, as well as the fraction of cells detectably superinfected as measured by reporter protein translation (**Fig. 4B-D, Fig. S3A**).

Inspection of the traces following single cells during superinfection revealed further quantitative characteristics of the exclusionary phenomenon (**Fig. 4E-F, Fig. S3B**). If infection by the founding virus progressively created a less favorable environment for the second virus, perhaps the single cell traces would reveal an effect on dynamics later during infection, such as a sudden shut-off of replication. Instead, superinfection replication was apparently inhibited at the earliest stages, as the max replication rate *m* of the second virus was reduced, as opposed to a late effect during replication (**Fig. S3C**).

To determine whether there might also be a bidirectional interaction between the two viruses, we plotted the final levels of mTurquoise2 and mCitrine at 12hpi across all temporal delays and superinfecting MOIs. We observed a striking anticorrelation between their endpoint fluorescence levels (**Fig. 4G, S3D**, Pearson correlation coefficient r=-0.99), suggesting that the superinfecting virus is equally able to reduce replication levels of the first virus, and that cells appear to have a fixed carrying capacity which determines the combined replication level of the two viruses.

Based on the observations of stereotypical growth dynamics, an early reduction in maximum production rates of the second virus during superinfection, and bidirectional interference during viral competition, we constructed a mathematical model of competitive Lotka-Volterra logistic growth using parameters estimated from single-virus infection experiments (**Fig. S3E, Methods**). This form predicts that, due to the speed of Sindbis replication, any temporal delay or decrease in the relative MOI would strongly disadvantage the second virus. The model recapitulates the empirical measurements, wherein we observed a graded response in superinfection replication levels with increased delays, decreased relative MOI, and combinations thereof leading to a corresponding reduction in the second viruses’ replication (**Fig. 4H**). Together, these data emphasize the importance of intrinsic growth kinetics in Sindbis superinfection exclusion.

## Discussion

While the complex choreography of polyprotein processing controlling the alphaviral lifecycle has been studied in detail, the early replicase activity, and overall trajectory of viral RNA synthesis *in situ* has remained unclear. In this study, we have sought to utilize single-cell imaging approaches in order to elucidate these phenomena, and to determine how these give rise to superinfection exclusion.

Current models of replication for alphaviruses have suggested that immediately following release of the polymerase nsP4 from the polyprotein P1234 by cis-cleavage, P123+nsP4 preferentially act as a minus-strand replicase. Reports have conflicted, however, regarding the activity of P123+nsP4 as a plus-strand replicase **(Lemm, Rümenapf et al. 1994, Shirako and Strauss 1994)**. The results shown here appear to rule out a scenario where the production of minus strand is first heavily favored during early infection, and instead suggests that polyprotein P123+nsP4 may additionally acts as a plus-strand replicase early during infection, yielding a replicase with a “balanced polarity”. This would be advantageous for the virus, as the early production of additional plus-strands would accelerate the onset of the exponential phase of replication, especially when such kinetics appear closely related to superinfection exclusion. Live-cell microscopy further confirmed that the speed of this onset was dependent on the MOI, while the maximum slope was not, as would be suggested by a model of logistic growth. These single-cell traces uncovered the sharpness of exponential growth and subsequent plateau, which would appear shallower and delayed when averaging over such an un-synchronized population.

A mechanism that provides a competitive advantage over a secondary similar infection within a fraction of an hour could be broadly beneficial across a wide host-range, especially considering the virus’ ability to enter a cell within mere minutes **(Helenius, Kartenbeck et al. 1980, Singh and Helenius 1992)**. Through the use of single-cell methods, we observe that the rapid onset of viral RNA synthesis with balanced polarity can confer such an advantage. Such a passive superinfection exclusion mechanism may also shield the infecting virus from any of its mutated progeny arising during naturally error-prone replication by the RNA-dependent RNA polymerase. Furthermore, a Lotka-Volterra model of exponential growth in a resource-limited environment appears consistent with empirical measurements of single and competitive viral replication, and independent of mutations ablating host-shutoff of transcription and translation **(Akhrymuk, Frolov et al. 2018)**. However, while these experiments suggest that a cell’s carrying capacity may be responsible for observed kinetics, the potential role of a “virus-intrinsic” program governing replication could also contribute to the logistic growth **(Zeng, Skinner et al. 2010, Razooky, Pai et al. 2015)**. Additionally, the apparent distribution in plateau levels of viral reporter fluorescence also highlights cell-to-cell differences and stochasticity in replication. The phenotypic state of cells and their spatial positions in culture have previously been shown to contribute to infection outcome **(Snijder, Sacher et al. 2009, Cohen and Kobiler 2016)**. Such factors, potentially including a range of characteristics such as cell-size, expression levels of key host-factors, innate-immune response, or transcriptional and translational capacity, can be considered together as an overall cellular state. Thus, the apparent “carrying-capacity” of each cell is therefore likely an amalgamation of some or all of these effects. The results described here, following previous studies of superinfection exclusion in alphavairuses have been performed in BHK cells, arguing against a role for innate immunity in this phenomenon. Indeed, should a host resource be the dominant component in limiting cellular capacity for alphaviral replication, the identification of such a target could potentially elucidate a novel drug target against alphaviral replication.

A comparative and integrative view of cell-state and innate immune activity in the context of viral replication, *in situ*, could reveal whether and how the observed dynamics may be further shaped by distinct immune responses between Sindbis virus’ mosquito and vertebrate hosts, and may also reveal differences in strategies used by the virus across these different organisms. As demonstrated here, single-cell imaging modalities can offer unique insight into classic questions of virology. As these methods are non-destructive, they are also suited to studying spatial properties of virus spreading in 2D monolayers, as well as more complex 3D environments like spheroids and organoids, potentially enabling an unprecedented look at the host-pathogen interplay through live cell reporters of innate immunity. Conversely, these powerful imaging strategies are equally effective at much shorter length-scales, for example, in measuring sub-cellular localization and trafficking. Thus, we anticipate that the further quantitative study of replication dynamics in complex interplay with innate immunity and stochasticity will be broadly relevant to the study of many infectious diseases.

## Methods

### Virus construction and production

Sindbis virus from Toto1101 **(Rice, Levis et al. 1987, Grakoui, Levis et al. 1989)** was transcribed *in vitro* using mMessage mMachine SP6 Transcription Kit (Ambion AM1340) for 2hrs, followed by DNAse Digestion for 20 minutes, and subsequently purified using RNEasy Mini Kit (Qiagen 74104). 500ng RNA was transfected into BHK-21s using Lipofectamine MessengerMax (Lifetech LMRNA001) into wells of a 24 well plate and passaged twice for expansion. Inoculations were performed for 20-30 minutes in 1%FBS in PBS with Calcium and Magnesium. Virus was concentrated using Amicon Ultra-15 Centfigiual Filter Concentrators (Millipore UFC910008). Virus was titered by plaque assay, and by fluorescence imaging to identify where ~50% of cells were infected for determination of MOI 1, 5hpi.

To construct reporter viruses, Toto1101 was digested with SpeI, and Gibson assembled with fluorescent proteins mCitrine and mTurquoise2 PCR’d with homology to surrounding the SpeI cut sites. Similarly, transcriptional and translational mutants in nsP2 and nsP3 were constructed serially using the same method.

### ImmunoFISH sample preparation and staining

Plus strand FISH probes were designed to bind over nsP1 and nsP2 sequences, which therefore exclude subgenomic RNA. Minus strand probes were targeted against the complement of nsP3 and nsP4 (**Table 1**). Cells were fixed in 4% formaldehyde for 5 minutes at room temperature, washed twice in PBS, and placed in 70% ethanol overnight at −20°C. The next morning, cells were washed twice with PBS; permeabilized in 0.1% TritonX for 5 minutes; washed twice in PBS; blocked in 10% Normal Goat Serum (Thermo 50062Z) treated with RNASecure (Invitrogen AM7005) for 30 minutes; washed twice with PBS; incubated with J2 primary-antibody (Scicons 10010200) at 0.5ug/ml in 10% Normal Goat Serum treated with RNASecure for 2 hours; washed twice with PBS; incubated with goat anti-Mouse Alexa Fluor 647 secondary antibody (Invitrogen A-21235) at RT for 30 minutes; washed twice in PBS and incubated with 10% Normal Goat Serum treated with RNASecure for an additional 10 minutes at RT; washed twice in PBS; postfixed in 4% formaldehyde for 5 minutes at RT; washed twice with PBS; equilibrated in FISH Wash Buffer containing 2X SSC (Invitrogen 15557044) and 20% Formamide (Ambion AM9342) for 5min at RT; and hybridized with Stellaris FISH probes labeled with Quasar 570 or Quasar 670 at 125nM (Biosearch Technologies, **Supplementary Table 1**) overnight at 30°C in Hybridization Buffer (containing 20% Formamide (Ambion AM9342), 2X SSC, 0.1g/ml Dextran Sulfate (Fisher Sci BP1585-100), 1mg/ml E.coli tRNA (Roche 10109541001), 2mM Vanadyl ribonucleoside complex (NEB S1402S), and 0.1% Tween 20 (VWR 97062-332) in nuclease free water). The next morning, the hybridization buffer was removed and cells were washed twice in FISH Wash Buffer; incubated in FISH Wash Buffer without probe for 30min at 30°C; washed three times with 2X SSC; counterstained with DAPI; and finally imaged in 2X SSC.

In the experiment to determine whether minus-strand probes could bind to dsRNA disrupted by sodium hydroxide, samples were washed twice in ddH_2_O, after fixing and PBS washing, then treated for 30s with 50mM NaOH and then washed three times in PBS. After this, the standard smFISH protocol was followed.

### Microscopy

Cells were imaged on a Nikon Ti2 with PFS4, a Nikon Motorized Encoded Stage, Lumencor SpectraX Light Engine, custom Semrock filters, and a Prime 95B sCMOS camera. Automated acquisition for snapshots and time-lapse was programmed in NIS Elements. The scope was equipped with an OKO stagetop incubator with temperature-, humidity-, and CO_2_-control, enabling long-term imaging. ImmunoFISH utilized a 60x objective, and time-lapse a 20X ELWD objective.

### Tissue Culture

BHK-21s (ATCC CCL-10) were grown at 37°C and 5% CO_2_ in DMEM/F12 with GlutaMax and Sodium Pyruvate (Gibco 10565018), supplemented with 10% FBS (Gibco 10437028, Lot 1780025), and NEAA (Gibco 11140050). For live-cell imaging of BHKs, DMEM/F12 was replaced with FluoroBrite DMEM (Gibco A1896701). For segmentation purposes for live cell imaging, BHKs were additionally stained with 1uM CellMask Red CMTPX cytoplasmic dye (Invitrogen C34552) for 10min at room temperature followed by three washes with PBS prior to imaging, and with 3ul/ml NucRed nuclear dye (Invitrogen R37106) or 5ul/ml NucBlue (Invitrogen R37605).

### Time-Lapse Image Analysis

All image- and data-analysis was performed in MATLAB 2019b using custom-built scripts. Single-cell tracking code was developed in the Elowitz Lab **(Hormoz, Singer et al. 2016)**, and modified for use with cytoplasmic cell segmentation. BHK’s which maintain large and irregular cytoplasmic boundaries require two markers, one nuclear and one cytoplasmic, for use in a seeded watershed segmentation algorithm. A movie of one well with no cells but all media and staining components is used to background correct sample wells at each corresponding time-point. Total fluorescence over the segmented area is summed to yield total fluorescence.

### ImmunoFISH Analysis

Two-channel segmentation is performed for immunoFISH using the same algorithm as in time-lapse imaging, but where autofluorescence on the 594 replaces the CellMask Red CMTPX dye as a cytoplasmic marker. Cells were imaged over a 6um z-stack every 2um; then a maximum-intensity projection was used for analysis. Puncta are identified using a Laplacian-of-Gaussian convolution as previously described **(Raj, Peskin et al. 2006, Singer, Yong et al. 2014)** on which a threshold is defined by an uninfected sample. Total fluorescence is analogously summed over the cell area.

### Modeling, Simulations, and Parameter Estimates

Logistic growth of the virus is modeled as

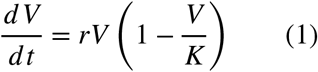

where parameters *r* and *K* are the replication rate and carrying capacity, respectively.

To estimate parameters from single infection experiments, we first subtract the initial fluorescence values and set the starting value equal to the MOI of the experiment. Then we rescale the median of the endpoints to be 200,000, or the number of copies of full length plus-strand previously estimated in the literature **(Wang, Sawicki et al. 1991)**. Finally, each trace is aligned so that its maximum slope occurs at the mean τ for the sample. These individual traces are then used in least squares curve fitting on the solution to equation 1, where *K* and *r* were allowed to vary, and the initial condition was fixed at the empirically determined MOI. On average, the slope over the last 3 hours was ~95% reduced from their maximum production rates (**Fig. 3H**). In fitting these traces to logistic growth, we selected only those whose final slope changed < 15% over the last three hours of the experiment (black lines in **Fig. S3B**; excluded traces are shown in gray). The estimated growth trace is shown as the heavy dashed line in **Figure S3B**.

Competitive Lotka-Volterra growth is modeled for virus 1 and 2 (V1 and V2, respectively) as:

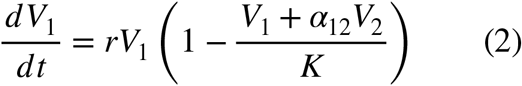

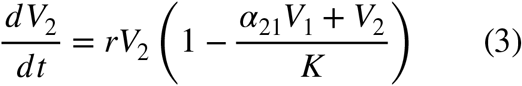

where the replication rate *r* is assumed to be the same for both viruses, and have a shared carrying capacity, *s*. Note that the coefficients α_12_=α_21_ = 1, a regime where there is no additional inhibitory effect of one virus on the either beyond their equal contribution towards their shared carrying capacity.

All modeling and estimation was performed in MATLAB 2019b.

## Data and Software Availability

Data and software are available upon request.

## Author Contributions

All authors conceived of and designed experiments together. Z.S.S. performed all experimental work and computational analysis. C.M.R. and T.D. supervised research. Z.S.S. wrote the manuscript with input from all authors.

## Acknowledgements

We are grateful to Professors Andrea Branch, William M Gelbart, and Sondra Schlesinger, and Dr. Brandon Razooky for providing critical feedback on the manuscript, as well as members of the Rice laboratory for regular discussions. This work was supported by National Institutes of Health NRSA award to Z.S.S, F32CA225145.

---

**Figure S1.**
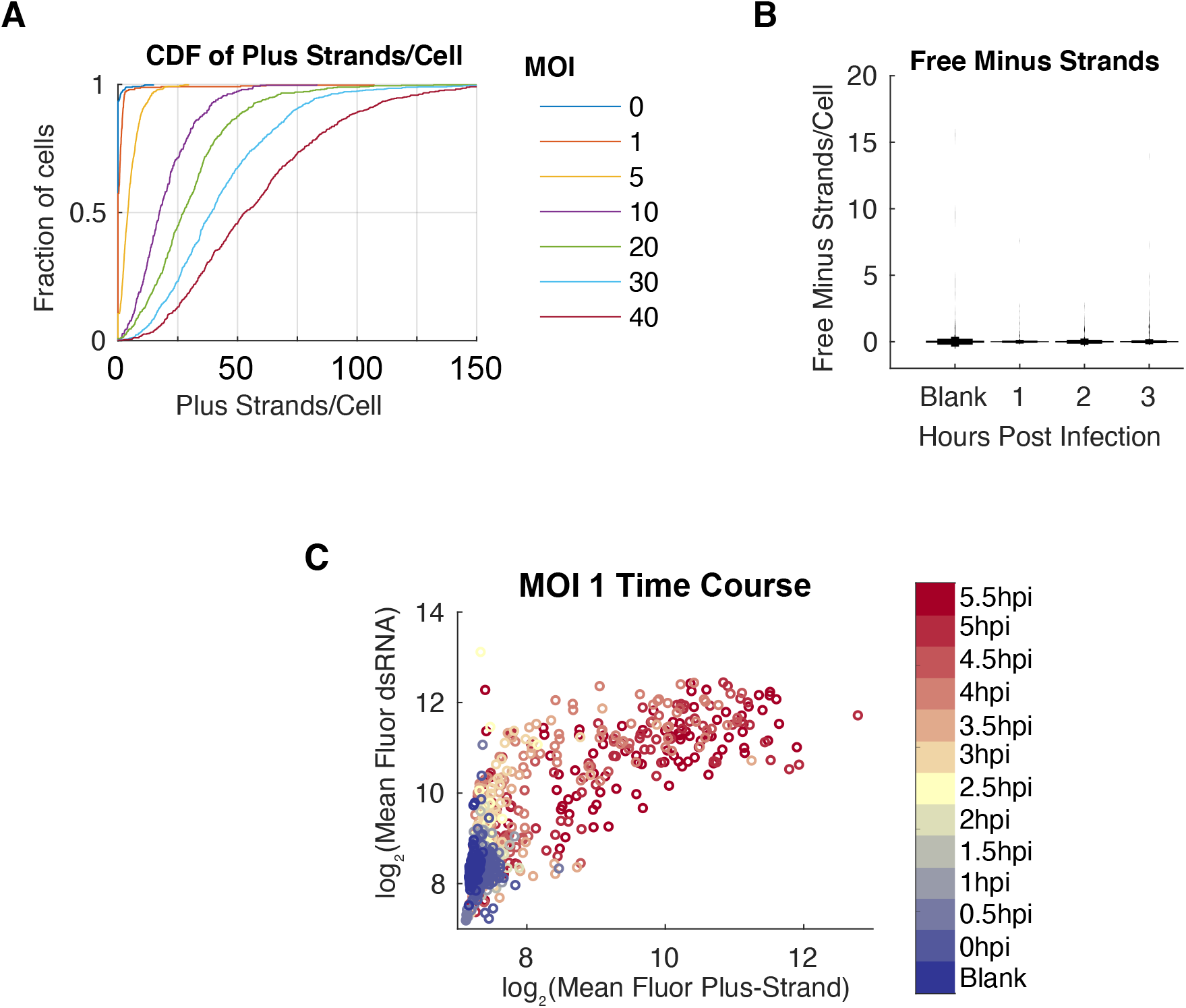
Supplementary Figure Corresponding to Figure 2. **A)** Cumulative distribution function of plus-strands per cell, immediately following inoculation. Data shown corresponds to Fig 2C. **B)** Detection of single molecules of free minus strand through 3 hours post infection. **C)** Time course of mean dsRNA fluorescence vs plus-strand mean FISH intensity every thirty minutes at MOI 1, analogous to Figure 2H.

**Figure S2.**
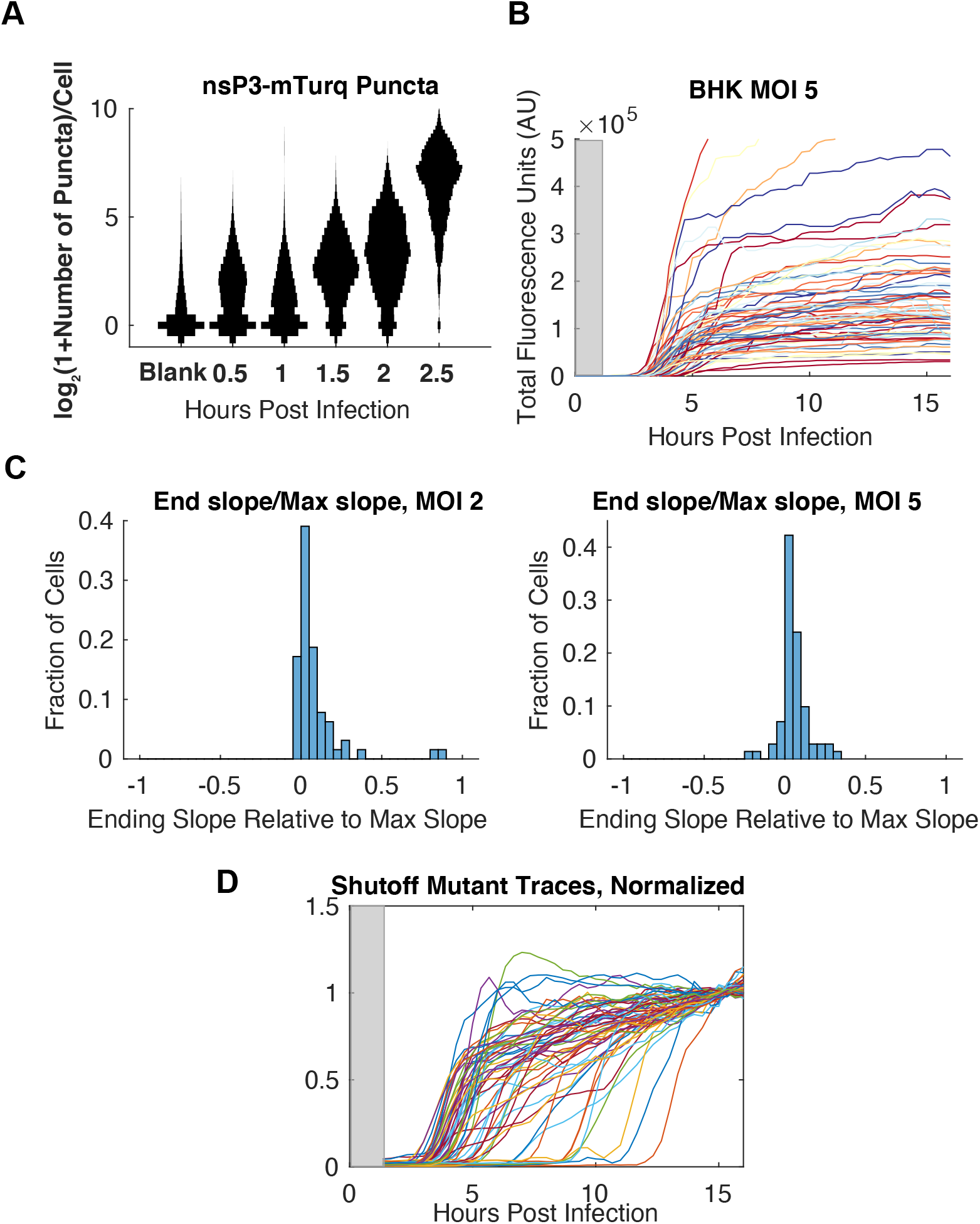
Supplemental Figure Corresponding to Figure 3. **A)** Counts of discrete spots in individual cells of nsP3-mTurquoise2 over a time course through 2.5h, displayed as smoothened vertical histograms. **B)** Single cell traces of total fluorescence infected at MOI 5. Gray box indicates the first hour during the inoculation and movie-preparation window before measurements are taken. **C)** The fold reduction in slopes computed as the slope over the last three hours of the traces fit to a line, divided by the maximum growth of the trace, shown for MOIs 2 and 5. **D)** Growth kinetics of nsP2-N24A/nsP3-P683Q double mutant.

**Figure S3.**
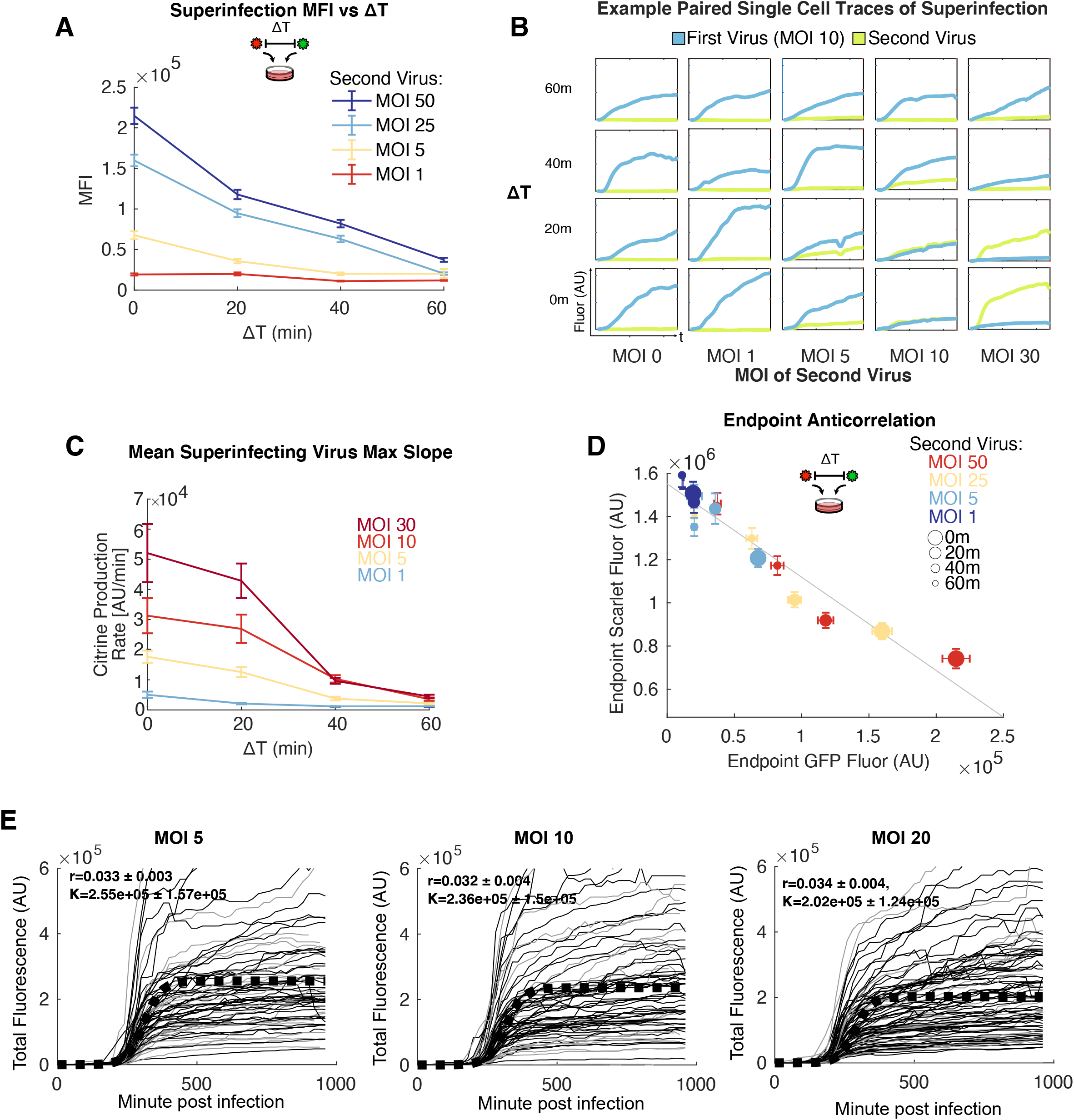
Supplemental Figure Corresponding to Figure 4. **A)** Here, SINV/nsP3-mScarlet is used as the first virus at MOI 5, followed by superinfection with SINV/nsP3-GFP at a range of MOIs and ΔTs, in order to rule out possible FRET. Analogous to Figure 4B, shown is fluorescence of the superinfecting virus per area (or MFI) averaged over all cells in the given condition as a function of delay (ΔT), shown as mean±SEM. Color coding of each line represents a different superinfecting MOI. **B)** Under superinfecting conditions, each panel shows an example pair of traces from both viral reporters in the same single cell, over time. To show both reporters on the same plot, y-axes for mCitrine and mTurquoise2 are distinct in a given plot, but the ranges are constant across all panels. MOIs indicated along the bottom are of the superinfecting SINV/nsP3-mCitrine, while the first virus, SINV/nsP3-mTurquoise2, is kept at MOI 10. Each row is a fixed delay between the first and second virus. Analogous to Fig 4E and F. **C)** The average maximum slope of the superinfecting virus as a function of the delay between the introduction of the first and second viruses. **D)** Here, SINV/nsP3-mScarlet is used as the first virus, followed by superinfection with SINV/ nsP3-GFP, in order to rule out possible FRET. Analogous to Figure 4G. Total fluorescence averaged over single-cells of nsP3-mScarlet versus nsP3-GFP at the end of the superinfection movie. Circle size represents delay between the first and second viruses, and color represents the MOI of the second virus, each shown at mean±SEM. Gray line is a linear-least squares fit, with a Pearson correlation coefficient of r=-0.99. **E)** Fits of logistic growth to individual traces across multiple MOIs. Black traces that increase less than 15% over the last three hours of the movie (black lines) are used for fitting, while the remainder are not (gray lines). See Methods.

**Table S1. Supplementary Table of All smFISH Probe Sequences**

**Supplemental Movie S1. Example time-lapse fluorescence movie of SINV/nsP3-mCitrine.** Infection performed at MOI 10. Red is the nuclear marker, cyan is the viral reporter fluorescence. Movie starts 1 hour post infection, with frames taken every 20min thereafter.

